# The molecular appearance of native TRPM7 channel complexes identified by high-resolution proteomics

**DOI:** 10.1101/2021.07.09.451738

**Authors:** Astrid Kollewe, Vladimir Chubanov, Fong Tsuen Tseung, Alexander Haupt, Catrin Swantje Müller, Wolfgang Bildl, Uwe Schulte, Annette Nicke, Bernd Fakler, Thomas Gudermann

**Affiliations:** Institute of Physiology II, Faculty of Medicine, University of Freiburg, Hermann-Herder-Str. 7, 79104 Freiburg, Germany; Walther-Straub Institute of Pharmacology and Toxicology, LMU Munich, Germany; Signalling Research Centres BIOSS and CIBSS, Freiburg, Germany; German Center for Lung Research, Munich, Germany.

**Keywords:** TRPM7, ARL15, CNNM, zinc, magnesium, calcium

## Abstract

The transient receptor potential melastatin-subfamily member 7 (TRPM7) is a ubiquitously expressed membrane protein consisting of ion channel and protein kinase domains. TRPM7 plays a fundamental role in the cellular uptake of divalent cations such as Zn^2+^, Mg^2+^ and Ca^2+^, and thus shapes cellular excitability, plasticity and metabolic activity. The molecular appearance and operation of TRPM7 channel complexes in native tissues have remained unresolved. Here, we investigated the subunit composition of endogenous TRPM7 channels in rodent brain by multi-epitope affinity purification and high-resolution quantitative MS analysis. We found that native TRPM7 channels are high molecular-weight multi-protein complexes that contain the putative metal transporter proteins CNNM1-4 and a small G-protein ARL15. Heterologous reconstitution experiments confirmed the formation of TRPM7/CNNM/ARL15 ternary complexes and indicated that ARL15 effectively and specifically impacts TRPM7 channel activity. These results open up new avenues towards a mechanistic understanding of the cellular regulation and function of TRPM7 channels.

**Impact Statement:** High-resolution proteomics in conjunction with biochemical and electrophysiological experiments revealed that the channel-kinase TRPM7 in the rodent brain forms macromolecular complexes containing the metal transporters CNNM1-4 and a small G protein ARL15.

## Introduction

TRPM7 encodes a bi-functional protein with a transient receptor potential (TRP) ion channel domain fused to a C-terminal α-type serine/threonine-protein kinase (reviewed in (1-3)). Among all other known channels and kinases, only its homologue TPRM6 shows a similar design (3, 4).

TRPM7 is involved in various cellular processes such as homeostatic balance, cell motility, proliferation, differentiation and regulation of immune responses (1-3). Genetic deletion of TRPM7 in mice is embryonically lethal, and tissue-specific null mutants have shown defects in cardiac and renal morphogenesis, organismal Zn^2+^, Mg^2+^, and Ca^2+^ homeostasis, thrombopoiesis, and mast cell degranulation (5-13). Besides, TRPM7 has emerged as a promising therapeutic target for numerous pathophysiological conditions (1-3, 14-16).

The channel-coding segment of TRPM7 comprises six transmembrane helices with a pore-loop sequence between S5 and S6 (Figure 1A, (17, 18)). Four subunits assemble to form constitutively active channels highly selective for divalent cations such as Zn^2+^, Ca^2+^, and Mg^2+^ (19-21). Free Mg^2+^, the Mg·ATP complex, and phosphatidylinositol-4,5-bisphosphate (PIP_2_) were described as physiological regulators of the channel activity of TRPM7 (19, 22). While Mg^2+^ or Mg·ATP act as negative regulators, PIP_2_ appears to be a crucial co-factor of the active channel (19, 22). Mechanistically, however, the effects of Mg^2+^, Mg·ATP, or PIP_2_ on TRPM7 activity are poorly understood, and most likely, there are additional regulators of TRPM7 function with hitherto unknown molecular identity.

**Figure 1.**
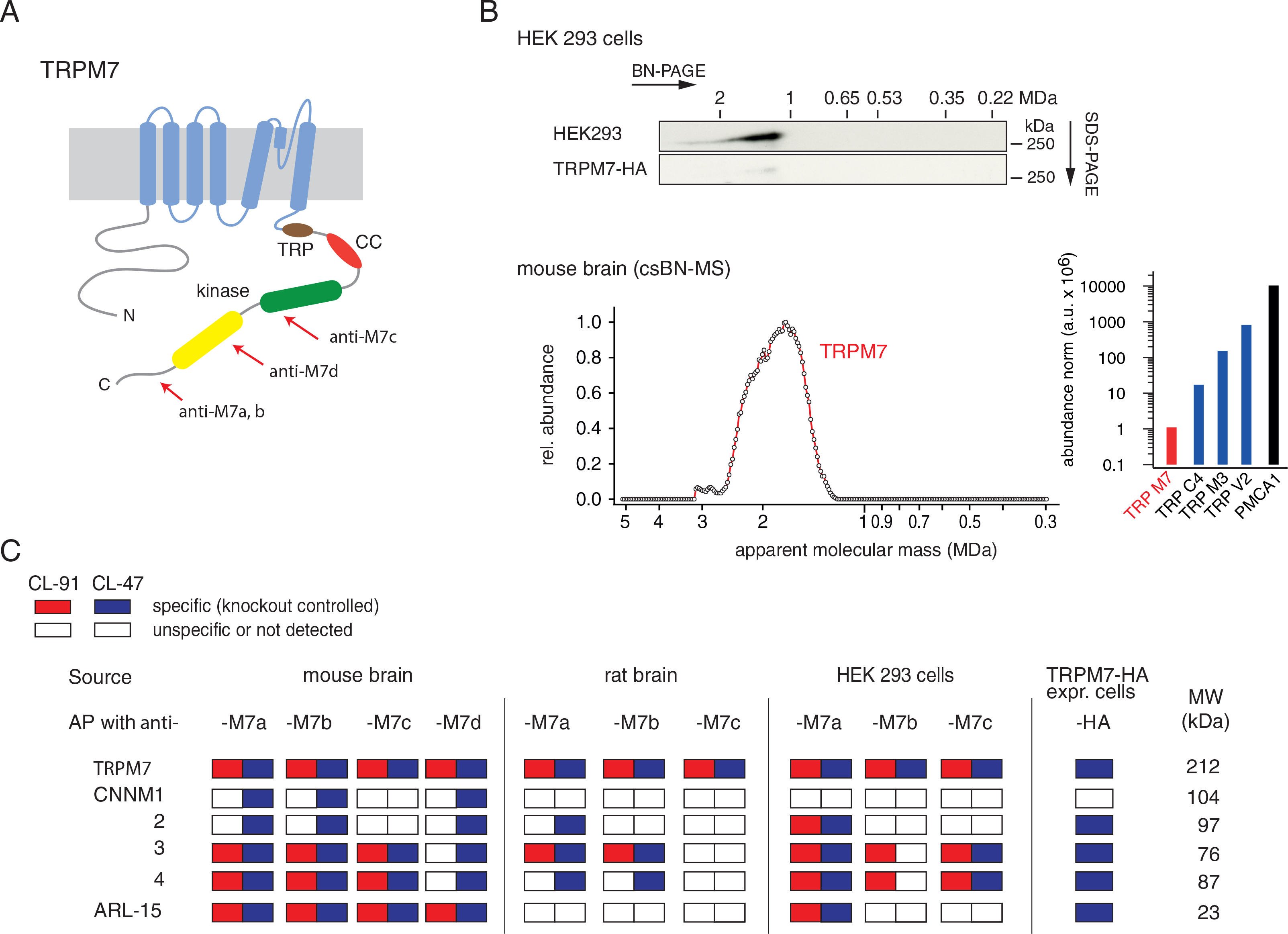
Protein constituents of native TRPM7 channels identified by multi epitope antibody-based affinity purification (ME-AP) proteomics. (**A**), Topology and localization of the *anti-TRPM7* antibodies used for ME-APs. Established hallmark domains of TRPM7 are colour-coded, TRP (transient receptor potential domain, brown), CC (coiled-coil domain, red), kinase (kinase domain, yellow), SD (serine/threonine-rich substrate domain of kinase(s), green). (**B**), Upper panel: Two-dimensional gel separation of TRPM7 channels in CL-47 solubilized membrane fractions of HEK293 cells with (upper panel) or without (lower panel) transfection of HA-tagged TRPM7, Western-probed with the indicated antibodies. Size (BN-PAGE) and molecular weight (SDS-PAGE) are as indicated, representative images of three experiments are shown. Lower panel: Abundance-mass profile of TRPM7 obtained by cryo-slicing BN-MS in a CL-47 solubilized membrane fraction from adult mouse brain (a total of 255 gel slices); inset: Abundance of the indicated proteins in the mouse brain. Note the large apparent molecular mass of the native TRPM7 channel in both culture cells and mouse brain, markedly exceeding the mass calculated for tetrameric channel assemblies (about 850 kDa, red circles). (**C**), Table summarizing the results of all *anti-TRPM7* APs performed with the indicated antibodies on membrane fractions prepared from rodent brain and cultured HEK293 cells. Colour coding is given in the upper left; MW is indicated on the right. TUC refers to series of APs with target-unrelated antibodies. Note that TRPM7 channels co-assemble with all CNNM family members and ARL-15 in the brain and epithelia. See also Figure 1-figure supplement 1; Supplementary file 1

The C-terminal alpha-kinase domain of TRPM7 acts in two ways: First, it auto-phosphorylates cytoplasmic residues of TRPM7, and second, it may target a variety of proteins with diverse cellular functions such as annexin A1, myosin II, eEF2-k, PLCγ2, STIM2, SMAD2, and RhoA (20, 23-28). In immune cells, the TRPM7 kinase domain has been reported to be clipped from the channel domain by caspases in response to Fas-receptor stimulation (29). In line with this observation, cleaved TRPM7 kinase was detected in several cell lines and shown to translocate to the nucleus, where it promotes histone phosphorylation (30).

The majority of the current knowledge about TRPM7 was derived from *in vitro* experiments with cultured cells, whereas insights into the operation of both channel and α-kinase activity of TRPM7 in native tissues are limited.

We, therefore, investigated the molecular architecture of TRPM7 in rodent brain by using blue native gel electrophoresis (BN-PAGE) and multi-epitope affinity purifications (ME-APs) in combination with high-resolution quantitative mass spectrometry (MS). These approaches showed that native TRPM7 channels are macromolecular complexes with an apparent size of ≧ 1.2 MDa and identified proteins CNNM1-4 and ARL15 as complex constituents. Subsequent functional studies in *Xenopus laevis* oocytes suggested ARL15 as a potent, hitherto unrecognized regulator of TRPM7 ion channel activity.

## Results

### ME-AP proteomic analyses of native TRPM7 channels

TRPM7 channels assemble from four subunits [2], each of which is about 1860 aa in length and comprises several distinct domains in its extended intracellular N- and C-termini in addition to its transmembrane channel domain (Figure 1A). Unexpectedly, analysis by native gel-electrophoresis (BN-PAGE) of TRPM7 channels either endogenous to HEK293 cells or exogenously expressed in these cells via transient transfection, elicited a molecular mass of at least 1.2 MDa considerably exceeding the molecular mass of ∼850 kDa calculated for TRPM7 tetramers (Figure 1B, upper panel). To see whether this large molecular size is a peculiarity of HEK293 cells, we recapitulated the analysis for TRPM7 channels expressed in mouse brain using a recently developed technique that combines BN-PAGE with cryo-slicing and quantitative mass spectrometry (csBN-MS, (31)). In this approach, membrane fractions prepared from the entire mouse brain and solubilized with the mild detergent buffer CL-47 (32-34) are first separated on a native gel, which is subsequently embedded and cut into 300 µm gel slices using a cryo-microtome. In a second step, the protein content of each slice is analysed individually by nanoflow liquid chromatography tandem mass spectrometry (nanoLC-MS/MS), providing information on both the identity and amount of any protein in each slice; noteworthy, protein amounts are determined with a dynamic range of up to four orders of magnitude (34-36). As illustrated in Figure 1B, lower panel, csBN-MS analysis of mouse brain membranes detected the TRPM7 protein with an apparent molecular mass between 1.2 and 2.6 MDa, comparable to the results obtained from HEK293 cells (Figure 1B, upper panel). Moreover, the determination of the total protein amount by signal integration over all slices showed that TRPM7 expression in the brain is rather low compared to other members of the TRP family of proteins. Thus, the abundance of TRPM7 is about one to three orders of magnitude below that obtained for TRPC4, TRPM3, or TRPV2 (Figure 1B, lower right).

Together, these results strongly suggest that native TRPM7 complexes exceed the predicted molecular size of bare tetrameric assemblies in different cellular environments suggesting that the rather simplistic view on the molecular make-up of native TRPM7 modules has to be revised.

To identify proteins that may co-assemble with TRPM7, we used affinity-purifications with multiple antibodies targeting distinct epitopes of the TRPM7 protein (Figure 1A, Figure 1-figure supplement 1) and evaluated the respective eluates of HEK293 cells and rodent brains by high-resolution quantitative MS analysis (ME-APs, (32-35)). HEK293 cells were selected because these cells are widely used for functional assessment of endogenous and overexpressed TRPM7. The brain was chosen since TRPM7 plays a critical role in neurological injuries and synaptic and cognitive functions (15, 37, 38). For these ME-APs, membrane fractions prepared either from whole brains of adult mice and rats or from WT HEK293 cells were solubilized with detergent buffers of mild (CL-47) or intermediate (CL-91) stringency (33-35). Also, TRPM7 was affinity-isolated from HEK293 cells transiently (over)-expressing C-terminally HA-tagged TRPM7 using an *anti*-HA antibody.

In all APs, TRPM7 could be reliably detected under both solubilisation conditions (Figure 1C) with MS-identified peptides covering a large percentage of the primary sequence of TRPM7 in samples from mouse brain as well as HEK293 cells (up to 77% and 98%, respectively) (Figure 2).

**Figure 2.**
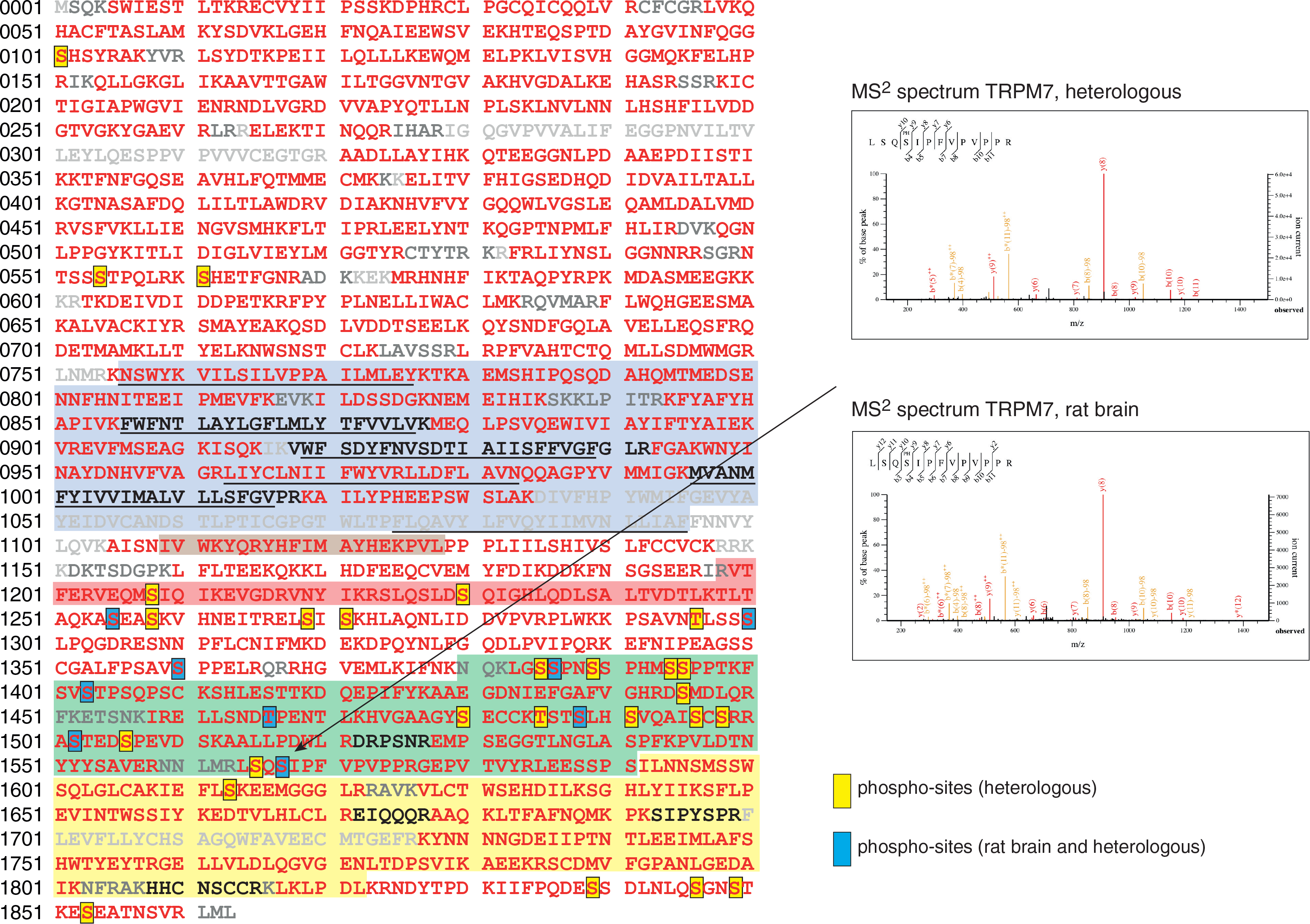
Coverage of the primary sequence of TRPM7 and phosphorylation sites as identified by MS-analyses in APs from transfected HEK293 cells and rodent brain. Peptides identified by mass spectrometry are in red; those accessible to but not identified in MS/MS analyses are in black, and peptides not accessible to MS/MS analyses used are given in grey. Blue boxes indicate Phospho-sites identified in the brain and transfected HEK293 cells; those uniquely seen in heterologous expressions are boxed in yellow. The MS/MS spectra on the right illustrate phosphorylation of Ser1567 in TRPM7 from both brain (upper) and culture cells (lower). Examples for all sites identified are given in supplementary file 3 to figure 2. Color-coding of hallmark domains is as in Figure 1A; S1-S6 helices of TRPM7 are underlined. See also Supplementary file 2 and Supplementary file 3

All other proteins identified in the ME-APs were evaluated for specificity and consistency of their co-purification with TRPM7 based on protein amounts determined by label-free quantification (see Methods section). The specificity of co-purification was assessed by comparing protein amounts in APs targeting TRPM7 with protein amounts obtained with stringent negative controls. Thus, (i) APs with four or five different target-unrelated control (TUC) antibodies were used as negative controls for *anti*-TRPM7 APs from rodent brain, (ii) *anti*-TRPM7 APs from a TRPM7^-/-^ HEK293 cell line (12) served as negative controls for *anti*-TRPM7 APs from WT HEK293 cells, and (iii) HEK293 cells heterologously expressing TRPM7-myc were used as negative controls for *anti*-HA APs from HEK293 cells overexpressing TRPM7-HA. A protein was considered consistently co-purified if detected in APs with at least two antibodies under the same solubilisation condition. Together, these consistency and specificity criteria identified five proteins as high-confidence interaction partners of TRPM7: the ADP-ribosylation factor-like protein 15 (ARL15) and the cyclin M family proteins CNNM1-4, putative Mg^2+^ transporters (Figure 1C, Table 1). Neither of these proteins was detected in any of the negative controls. Moreover, they were not only consistently co-purified with several antibodies but with the exception of CNNM1 also from both rodent brain and HEK293 cells. Comparison of the degree of association under the two solubilisation conditions revealed that the interaction between TRPM7, ARL15 and CNNMs was weakened by the more stringent detergent CL-91 (Figure 1C, Table 1).

**Table 1.** Protein constituents of native TRPM7 channels identified in ME-APs. Accession numbers refer to the UniProtKB database; colour-coding is given at the bottom. Relative abundance refers to the amount of TRPM7 as a reference and was classified as follows: = when between 0.33-fold and 3.3-fold of reference, < when between 0.033-fold and 0.33-fold of reference, << when between 0.0033-fold and 0.033-fold of reference, and <<< when less than 0.0033-fold of the reference amount. ^#^ co-purified with *anti*-M7a from HEK293 cells (CL-47) and with *anti*-M7c HEK293 cells (CL-91); ^##^ co-purified with anti-M7c from rat brain membranes (CL-91); ^###^ co-purified with anti-M7a from HEK293 cells (CL-47, CL-91). See also Supplementary file 1

| Protein ID | Acc. No.<br>UniProtKB | Name | primary<br>function | rel. abundance<br>CL-47 CL-91 |  |
| --- | --- | --- | --- | --- | --- |
| TRPM7 | Q923J1 | TRP channel M7 | ion channel | = | = |
| CNNM1 | Q0GA42 | Transporter CNNM1, Cyclin-M1 | pot. transporter | << |  |
| CNNM2 | Q5U2P1 | Transporter CNNM2, Cyclin-M2 | pot. transporter | < | << |
| CNNM3 | Q32NY4 | Transporter CNNM3, Cyclin-M3 | pot. transporter | < | << |
| CNNM4 | Q69ZF7 | Transporter CNNM4, Cyclin-M4 | pot. transporter | < | << |
| ARL15 | Q8BGR6 | ADP-ribosylation factor-like protein 15 | unknown | = | << |
| TP4A1# | Q93096 | Protein tyrosine phosphatase type IVA 1 | enzyme | <<< | << |
| TP4A3## | Q9D658 | Protein tyrosine phosphatase type IVA 3 | enzyme |  | << |
| TRPM6### | Q9BX84 | TRP channel M6 | ion channel | <<< | << |

■ transmembrane ■ cytoplasmic

In addition to subunit assembly, the MS-data provided further insight into the post-translational modification(s) of the TRPM7 protein. Thus, TRPM7 purified either from rodent brain or from transfected HEK293 cells showed very similar patterns of serine and threonine phosphorylation, reflected by matching MS/MS spectra of peptides harbouring phosphorylation sites (Figure 2). Out of the nine common phospho-sites, four have not been reported for TRPM7 in native tissue before (S1300, S1360, T1466, and S1567; Supplementary file 2 to Figure 2). An additional 26 phosphorylated serine and threonine residues could be assigned to TRPM7 isolated from HEK 293 cells, presumably based on the higher amounts of TRPM7 available for analysis from heterologous (over)-expression material. 22 of these 26 sites match sites previously reported for TRPM7 endogenously or heterologously expressed in cell lines, and four sites were newly detected (S1208, S1480, S1496, S1853; Supplementary file 2 to Figure 2). Most of the identified phosphorylation sites were found to cluster within the C-terminal cytoplasmic domain of TRPM7.

Next, we verified the identified interactions between TRPM7, ARL15 and CNNM1-4 in co-expression experiments performed in TRPM7^-/-^ HEK293 cells (Figure 3). Flag-tagged ARL15 and CNNM proteins could be specifically and robustly co-purified with HA-tagged TRPM7 in *anti*-HA APs when all three proteins were present, whereas the association was markedly less efficient when ARL15-Flag or CNNM-Flag were co-expressed with TRPM7-HA alone (Figure 3, Figure 3-figure supplement 1). These results corroborated the ME-AP results from the rodent brain and strongly suggested the formation of ternary complexes containing TRPM7, ARL15 and CNNM proteins.

**Figure 3.**
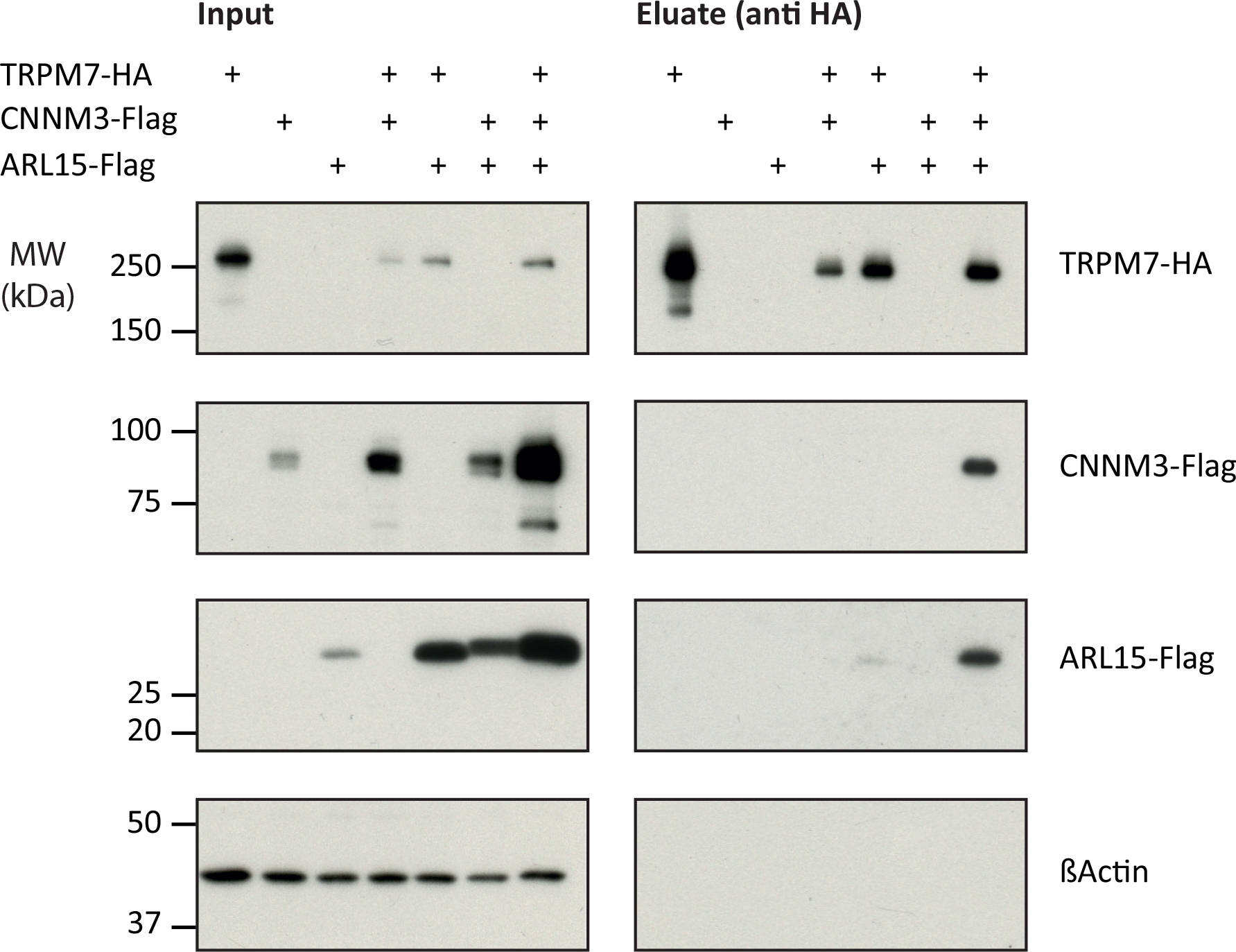
Heterologous reconstitution of TRPM7 complexes in HEK293 cells. APs with *anti*-HA antibody from CL-47 solubilised membrane fractions of *TRPM7* -/- HEK293 cells transiently expressing the proteins indicated above. Input and eluates of the distinct APs were separated by SDS-PAGE and Western-probed with *anti*-Flag and *anti*-HA antibodies. MW is marked on the left. Typical examples of three independent experiments. See also Figure 3-figure supplement 1.

### Effects of CNNM3 and ARL15 on TRPM7 channel activity

To investigate if the assembly of TRPM7 with ARL15 and CNNM proteins modified TRPM7 function, we studied their effect(s) on TRPM7 currents by co-expression in *Xenopus laevis* oocytes. This approach allows co-expression of defined protein ratios by cRNA injection and, therefore, is widely used for functional assessment of ion channel complexes, including functional interaction of TRPM7 with TRPM6 (6). The two-electrode voltage clamp (TEVC) measurement in Figure 4A illustrates a typical current-voltage (I-V) relationship of constitutively active TRPM7 channels characterized by steep outward-rectification and a reversal potential of around 0 mV (19). Co-expression of TRPM7 and CNNM3, the most efficiently co-purified CNNM protein (Figure 1C), neither changed the shape of the I-V relationship nor current amplitudes. In contrast, ARL15 effectively suppressed constitutive TRPM7 currents in a concentration-dependent manner, as deduced from experiments with increasing cRNA amounts (Figure 4B). The suppressive effect was specific for TRPM7, as co-expressed ARL15 did not inhibit another TRP channel, TRPV1, in an analogous experiment (Figure 4-figure supplement 1). Oocytes co-expressing all three proteins TRPM7, CNNM3, and ARL15, did not exhibit TRPM7 currents, similar to the co-expression of TRPM7 and ARL15.

**Figure 4.**
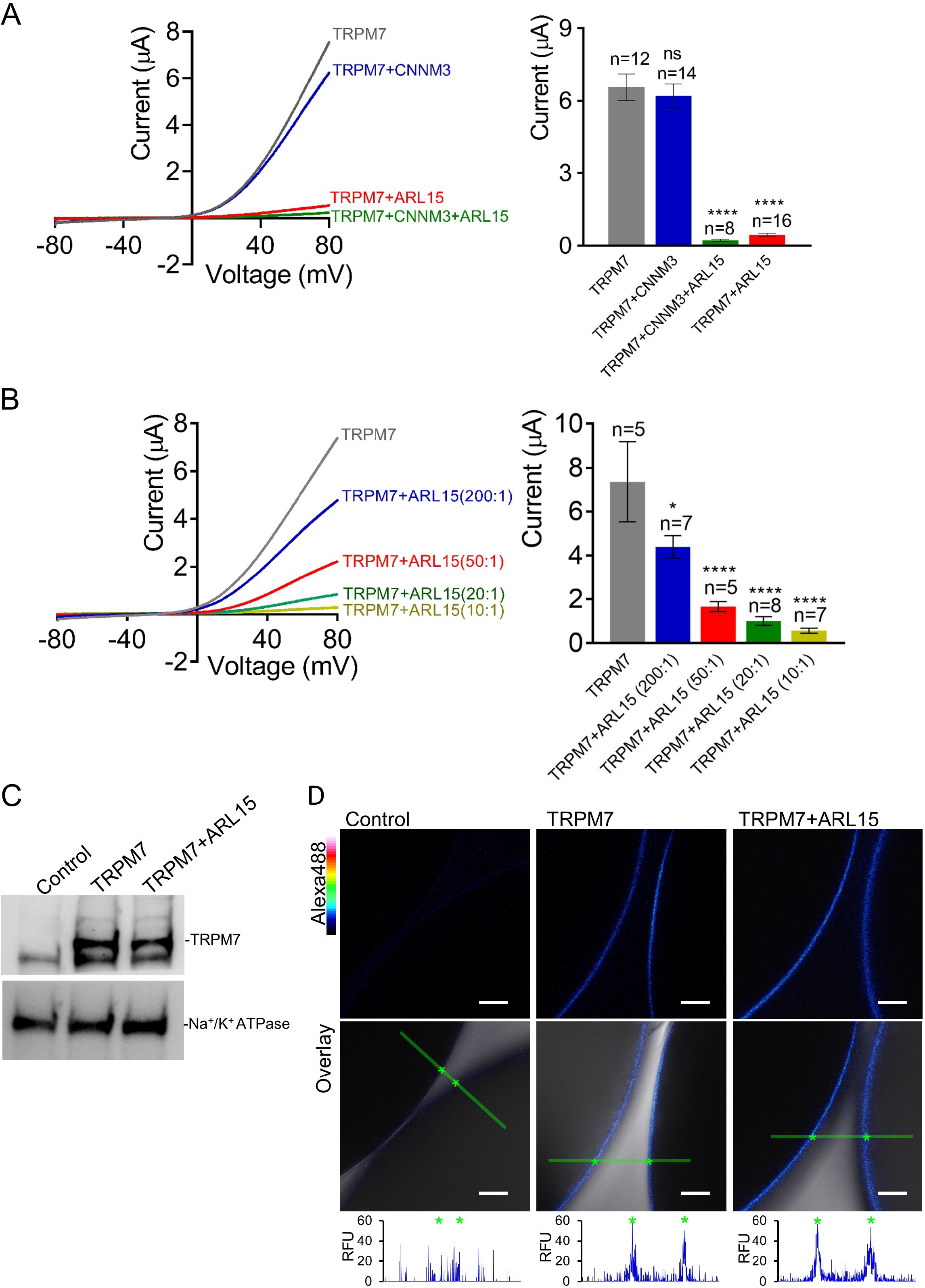
Heterologous expression of TRPM7 in *Xenopus* oocytes. **(A, B)** Two-electrode voltage clamp (TEVC) measurements of TRPM7 currents. (A) *Left panel*: Representative current-voltage (I-V) relationships of TRPM7 currents measured in oocytes expressing TRPM7 alone or TRPM7 with CNNM3 or ARL15 (cRNAs ratio 2:1), and TRPM7 with CNNM3 and ARL15 (cRNAs ratio 2:1:1). *Right panel*: Current amplitudes (mean ± SEM) at +80 mV in measurements shown on the left. Two independent batches of injected oocytes (n=8-16) were examined; * P < 0.05; **** P < 0.0001 (ANOVA). (B) *Left panel*: Representative I-V relationships of TRPM7 currents measured in oocytes expressing TRPM7 or co-expressing TRPM7 with ARL15 at the indicated ratios of injected cRNAs. *Right panel*: Current amplitudes (mean ± SEM) at +80 mV in measurements shown on the left. Two independent batches of injected oocytes (n=5-7) were examined; **** P < 0.0001 (ANOVA). (C) Western blot analysis of TRPM7 expression using *anti-*M7d in total lysates of oocytes injected with TRPM7 or TRPM7 and ARL15 cRNAs (ratio 10:1). Representative results are shown for two independent experiments. (D) Immunofluorescence staining of un-injected oocytes (Control) or oocytes injected with TRPM7 (TRPM7) or TRPM7 and ARL15 cRNAs (TRPM7+ARL15, ratio 10:1) using *anti-*M7d antibody and *anti*-mouse antibody conjugated with Alexa Fluor 488. Confocal images of Alexa Fluor 488 fluorescence (Alexa488) and overlays of Alexa488 with differential interference contrast images (Overlay) are depicted for two independent oocytes per image; scale bars, 50 μm. The rainbow indicator outlines fluorescence intensity. The diagrams depict fluorescence intensity acquired along the green bars shown in *Overlay* images. The stars indicate the cell surface of two oocytes. Typical examples of two independent experiments (n=10 oocytes) are shown. See also Figure 4-figure supplement 1.

Next, we examined if the interference of ARL15 with TRPM7 function was due to reduced expression levels or altered membrane localization. Western-blot analysis of oocytes injected with TRPM7 or TRPM7 and ARL15 cRNAs did not reveal any change in the expression level of TRPM7 (Figure 4C). Using immunofluorescent staining with the *anti-*M7d antibody, we detected TRPM7 at the cell surface of oocytes injected with TRPM7 but not in uninjected oocytes (Figure 4D). Notably, the TRPM7 signal was similarly detectable at the cell surface of oocytes co-expressing TRPM7 and ARL15 (Figure 4D).

Collectively, these results suggest that the inhibitory effect of ARL15 on TRPM7 currents is specific, dose-dependent and unlikely to be associated with reduced abundance of the channel at the cell surface; whether this modulatory effect is relevant for TRPM7 channel complexes in the rodent brain still remains to be resolved.

## Discussion

In the present study, we investigated the molecular appearance and subunit composition of TRPM7 as present in the cell membrane(s) of the rodent brain. We show that TRPM7 forms macromolecular complexes by assembling with CNNM proteins 1-4 and ARL15. Moreover, functional expression in a heterologous expression system showed that ARL15 might strongly affect TRPM7 channel function.

BN-PAGE of membrane fractions isolated from rodent brain and cultured HEK 293 cells identified endogenous TRPM7 in high ∼1.2 MDa molecular weight complexes exceeding the calculated molecular mass of TRPM7 tetramers (∼850 kDa) and suggesting that the TRPM7 channel-kinase is predominantly embedded in a large macromolecular complex.

Compared to other native TRP channels, such as TRPC4, TRPM3 and TRPV2, the expression level of TRPM7 was found to be up to three orders of magnitude lower, thus classifying TRPM7 as a very low-abundant protein in the rodent brain and indicating that comprehensive determination of the TRPM7 complexome is technically challenging. The unbiased ME-AP approach paired with stringent negative controls nevertheless allowed the identification of high-confidence interaction partners based on their specific and consistent co-purification with TRPM7. Consequently, five proteins were found to assemble with native TRPM7, including four members of the CNNM gene family encoding putative Mg^2+^ transporters CNNM1-4 and a small G-protein ARL15. The fact that we did not detect all the interactors seen in mouse brain also in APs from rat brain is most likely due to the low abundance of endogenous TRPM7 (∼50% less TRPM7 compared to APs from mouse brain).

Importantly, the interaction of TRPM7 with ARL15 and CNNM proteins was successfully confirmed in heterologous expression experiments (Figure 3). We also noted that previous proteome-wide interactome screens in cultured cells suggested an association of ARL15 with TRPM7 and CNNMs ((39), bioRxiv doi:10.1101/2020.01.19.905109), in line with our results. To obtain the first insight into a possible functional impact of ARL15 and CNNM3, the most prominent interaction partners of TRPM7 in our experimental settings, we measured the channel activity of mouse TRPM7 expressed in *Xenopus* oocytes. We found that co-expression of CNNM3 with TRPM7 did not lead to apparent changes in TRPM7 currents, whereas the addition of ARL15 caused complete suppression of TRPM7 activity, suggesting that ARL15 might be a critical regulatory factor of the TRPM7 channel.

The CNNM (cyclin M; CorC) gene family encodes highly conserved metal transporter proteins identified in all branches of living organisms, ranging from prokaryotes to humans (40, 41). There are four family members in mammals, CNNM1– CNNM4, widely expressed in the body and abundantly present in the brain (40, 41). Functional expression studies proposed that CNNMs operate as Na^+^/Mg^2+^ exchangers responsible for the efflux of cytosolic Mg^2+^ from the cell (40, 41). Consistently, genetic inactivation of CNNM4 in mice leads to systemic Mg^2+^ deficiency (42). In humans, point mutations in *CNNM2* cause hypomagnesemia (43), while mutations in *CNNM4* are associated with Jalili syndrome (44). Recently resolved crystal structures of two prokaryotic CNNM-like proteins revealed that CNNMs form dimers and that each monomer contains three transmembrane helices harbouring Mg^2+^ and Na^+^ binding sites consistent with the suggested Na^+^-coupled Mg^2+^ transport function of CNNMs ((45), bioRxiv doi.org/10.1101/2021.02.11.430706). While the majority of CNNM proteins in a cell is not bound to TRPM7, the direct association identified in this study suggests a new concept implying that two transporting mechanisms, TRPM7-mediated influx of divalent cations (Zn^2+^, Mg^2+^ and Ca^2+^) and CNNM-dependent Na^+^/Mg^2+^ exchange, can be physically coupled under native conditions, thus, warranting future studies to examine the exact functional interplay between TRPM7 and CNNMs.

ARL15 is a member of the ARF gene family of small G-proteins (46). A common feature of ARFs is their ability to bind and regulate effector proteins in a GTP-dependent manner (46). GDP- and GTP-bound states of ARFs are controlled by GTPase-activating proteins (GAP) in conjunction with GTP exchange factors (GEF) (46). The best-characterised ARFs are involved in membrane trafficking, phospholipid metabolism and remodelling of the cytoskeleton (46). While genome-wide association studies have linked ARL15 to systemic Mg^2+^ homeostasis and energy metabolism in humans (47, 48), the particular functional role and corresponding GAP, GEF and effector proteins of ARL15 remain to be established. To this end, the strong effect of ARL15 in suppressing TRPM7 currents observed in our study may suggest that TRPM7 serves as a specific effector protein of ARL15. The significance of this modulatory effect for native TRPM7 in the rodent brain, however, remains to be shown.

In some TRPM7-APs from HEK293 cells, we detected TRPM6, a genetically related channel, and two proteins representing the gene family of Phosphatase of Regenerating Liver 1 and 3 (PRL1, 3; also entitled Protein tyrosine phosphatases type 4A1 and 3, PTP4A1 and 3) (Table 1). The Mg^2+^ transporter protein TRPM6 has been described to physically and functionally interact with TRPM7 (6, 49, 50). In the present study, TRPM6, even though detected, could not be consistently co-purified with multiple anti TRPM7 antibodies, likely because TRPM6 is expressed at very low levels in the brain and HEK293 cells. Nevertheless, a previous study reporting that heterologously expressed ARL15 positively modulates TRPM6 (47) might suggest an overlap between the TRPM6 and TRPM7 interactomes.

Interestingly, a recent interactome assessment of HEK293 and HTC116 cells revealed that TP4A1 and TP4A2 also interact with ARL15 and CNNMs ((39), bioRxiv doi:10.1101/2020.01.19.905109). Furthermore, a hypothesis-driven search for interaction partners of CNNMs has shown that TP4A proteins assemble with CNNMs and that such interactions shape Mg^2+^ efflux from cells (51-56). These findings commensurate with our observation that TP4A1 and TP4A3 could be found in TRPM7 APs at low amounts.

Hence, based on the present analysis of native TRPM7 complexes in conjunction with earlier interactome experiments and functional expression studies, it is tempting to speculate that TRPM7/ARL15/CNNMs/TP4As form a protein network orchestrating transport of divalent cations across the cell membrane.

## Material and Methods

### Antibodies

Antibodies used for APs were: *anti*-HA (11867423001, Roche) and *anti*-HA (26183, Invitrogen). Target unrelated control (TUC) antibodies were: rabbit IgG (12-370, Millipore), *anti*-ßArrestin 2 (sc-13140, Santa Cruz), anti-TRPC1 (4921, a gift from Veit Flockerzi), *anti*-Sac1 (ABFrontier), *anti*-TRPC3 (1378, a gift from Veit Flockerzi), *anti*-NMDAR1 (MAB1586, Sigma), *anti*-LRRTM2 (23094-1-AP, ProteinTech), *anti*-DPP10 (sc-398108, Santa Cruz), and *anti*-RGS9 (sc-8143, Santa Cruz).

*Anti*-TRPM7 mouse monoclonal antibody (anti-M7a) was purchased from Thermo Fisher Scientific (clone S74-25, Product # MA5-27620). *Anti*-TRPM7 mouse monoclonal antibody (*anti*-M7b) was obtained from NeuroMab (clone N74/25, Product # 75-114). Generation of a rabbit polyclonal anti-TRPM7 antibody (*anti*-M7c) was described previously (26). Briefly, rabbits were immunised with a phosphorylated peptide H2N-DSPEVD(p)<u>S</u>KAALLPC-NH2 coupled via its C-terminal cysteine residue to keyhole limpet hemocyanin (Eurogentec, Belgium). The generated serum was subjected to two rounds of affinity chromatography: a fraction of the antibody was purified using the phosphorylated peptide followed by an additional round of chromatography using a non-phosphorylated variant of the peptide (H2N-DSPEVDSKAALLPC-NH2). The latter fraction of antibody was used in the present study.

Anti-TRPM7 2C7 mouse monoclonal antibody (*anti*-M7d, Figure 1-figure supplement 1) was produced by Eurogentec (Belgium) as follows. The nucleotide sequence coding for His_6_-tag followed by a cleavage site sequence for TEV protease and the amino acids 1501-1863 (kinase domain, KD) of mouse TRPM7 protein was synthesised *in vitro* and cloned into the prokaryotic expression vector pT7. The resulting expression construct pT7-His_6_-mTRPM7-KD was verified by sequencing and transformed in *E. coli* (BL21 DE3 pLysS). Next, the transformed *E. coli* strain was amplified in LB medium at 25°C. 1 mM IPTG was used for induction of the His_6_-mTRPM7-KD protein expression. The harvested cell pellet was disrupted by sonication. His_6_-mTRPM7-KD was identified in the soluble fraction of the lysate. His_6_-mTRPM7 was purified on a Ni Sepharose™ 6 Fast Flow column on an AKTA™ Avant 25 (GE-Healthcare) using an imidazole gradient of 20-500 mM. The fraction containing His_6_-mTRPM7-KD was dialysed against a Tris buffer (0.5 mM EDTA, 1mM DTT and 50 mM Tris HCl pH 7.5). His_6_-mTRPM7-KD was subjected to TEV protease (New England Biolabs) digestion according to the manufacturer’s instructions. Subsequently, non-digested His_6_-mTRPM7-KD and His_6_-tagged fragments were removed using a Ni-Sepharose^TM^ 6 Fast Flow column. The flow-through containing the cleaved mTRPM7-KD was concentrated to 0.5 mg/ml in the Tris buffer and stored at -80°C. SDS-PAGE was used to verify the removal of the His_6_-tag.

The standard mouse monoclonal antibody production program of Eurogentec (Belgium) was conducted to immunise four mice using the mTRPM7-KD protein and to produce a library of hybridomas. ELISA and Western-blot were used to screen the hybridomas and to perform a clonal selection. Two hybridoma clones, 2C7 and 4F9 (isotypes G1;K), were selected based on the antibody quality released in the culture medium. Both clones were propagated, and the corresponding cell culture media were collected for large-scale purification of the IgG fraction using Protein G affinity chromatography. The IgG fractions from 2C7 and 4F9 were dialysed in PBS and stored at -80 °C. The specificity of the 2C7 and 4F9 clones were verified by Western blot analysis of HEK293 cells overexpressing the TRPM6 and TRPM7 proteins (Figure 1-figure supplement 1). The 2C7 antibody detected the mouse or human TRPM7, but not the mouse or human TRPM6 (Figure 1-figure supplement 1). In contrast, the 4F9 antibody detected only the mouse TRPM7 (Figure 1-figure supplement 1). Consequently, the 2C7 antibody (*anti*-M7d) was used in the present study.

### Cell culture

HEK293 and HEK293T cells (Sigma) were cultured at 37 °C, 5% CO_2_ in Dulbecco’s Modified Eagle’s high glucose GlutaMAX medium (Gibco) supplemented with 10% fetal calf serum (Gibco), 1% penicillin/ streptomycin (Gibco) and 10 mM Hepes (Gibco). TRPM7^-/-^ HEK293T cells (12) were cultured as WT cells with an addition of 10 mM MgCl2, 3 µg/ml Blasticidin S (InvivoGen) and 0.5 µg/ml Puromycin (Gibco) to the medium.

### Molecular biology

Expression constructs encoding mouse TRPM7 and TRPM6 and human TRPM6 proteins (in pIRES2-EGFP vector) were reported previously (6, 49). The mouse TRPM7 cDNA in the pOG1 vector was generated as explained before (6). The mouse TRPM7-Myc and TRPM7-HA cDNA variants in pcDNA3.1/V5-His TA-TOPO vector were described earlier (6, 49). Expression constructs encoding Myc-Flag-tagged (C-end) mouse CNNM1-4 and ARL15 in the pCMV6-Entry expression vector were acquired from OriGene (MR218318 for CNNM1, MR218370 for CNNM2, MR224758 for CNNM3, MR215721 for CNNM4, MR218657 for ARL15) and verified by sequencing.

### Biochemistry

#### Transient transfection of cultured cells

WT HEK293T cells were transfected with polyethylenimine (Polysciences) using a DNA to polyethylenimine ratio of 1:2.5. For transfection of TRPM7^-/-^ HEK293T cells (12), plasmid DNA was diluted to 30 µg/ml in Hank’s balanced salt solution, precipitated by addition of 113 mM CaCl2 (final concentration) and added to the cells in culture medium lacking Blasticidin S, Puromycin and 10 mM MgCl2. For transfection, *TRPM7*, *ARL15* and *CNNM* plasmid DNAs were mixed at a ratio of 3:1:1.

#### Preparation of plasma membrane-enriched protein fractions

Freshly excised brains from 25 male and 25 female adult rats or mice were homogenised in homogenisation buffer (320 mM sucrose, 10 mM Tris/HCl pH 7.4, 1.5 mM MgCl2, 1 mM EGTA and protease inhibitors (Leupeptin (Sigma), Pepstatin A (Sigma), Aprotinin (Roth) (1 µg/ml each), 1 mM Phenylmethylsulfonyl fluoride (Roth), 1 mM Iodoacetamide (Sigma)), particulates removed by centrifugation at 1,080xg and homogenised material collected for 10 min at 200,000xg. After hypotonic lysis in 5 mM Tris/HCl pH 7.4 with protease inhibitors for 35 min on ice, the lysate was layered on top of a 0.5 and 1.3 M sucrose step-gradient in 10 mM Tris/HCl pH 7.4, 1 mM EDTA/EGTA, and the plasma membrane-enriched fraction collected after centrifugation (45 min, 123,000xg) at the interface. Membranes were diluted in 20 mM Tris/HCl pH 7.4, collected by centrifugation (20 min, 200,000xg), and resuspended in 20 mM Tris/HCl pH7.4.

Cultured cells were harvested in phosphate buffer saline with protease inhibitors, collected by centrifugation (10 min, 500xg) and resuspended in homogenisation buffer. After sonication (2x 5 pulses, duty 50, output 2 (Branson Sonifier 250)), membranes were pelleted for 20 min at 125,000xg and resuspended in 20 mM Tris/HCl pH 7.4. Protein concentration was determined with the Bio-Rad Protein Assay kit according to the manufacturer’s instructions.

#### Immunoprecipitation

Membranes were resuspended in ComplexioLyte CL-47 or CL-91 solubilisation buffer (Logopharm) with added 1 mM EDTA/EGTA and protease inhibitors at a protein to a detergent ratio of 1:8 and incubated for 30 min on ice. Solubilised protein was cleared by centrifugation (10 min, 125000xg, 4°C) and incubated with antibodies cross-linked to Dynabeads (Invitrogen) by overhead rotation for 2 h on ice. After two short washing steps with ComplexioLyte CL-47 dilution buffer (Logopharm), the captured protein was eluted in Laemmli buffer with dithiothreitol added after elution. Eluted proteins were separated by SDS-PAGE. For MS/MS analysis silver-stained (57) protein lanes were cut-out, split at 50 kDa and pieces individually subjected to standard in-gel tryptic digestion (58). For chemiluminescent detection, proteins were western blotted onto PVDF membranes and probed with the following antibodies: *anti*-HA (11867423001, Roche), *anti*-Flag (F3165, Sigma), *anti*- ßActin (bs-0061R, Bioss Inc.).

#### Blue-native Polyacrylamide gel electrophoresis

Two-dimensional BN-PAGE/SDS-PAGE protein analysis was performed as described previously (59). Membrane protein fractions were solubilised in ComplexioLyte CL-47 as described above, salts exchanged for aminocaproic acid by centrifugation through a sucrose gradient and samples loaded on non-denaturing 1-13% linear polyacrylamide gradient gels (anode buffer: 50 mM Bis-Tris, cathode buffer: 50 mM Tricine, 15 mM Bis-Tris, 0.02% Coomassie Blue G-250). For separation in the second dimension, individual gel lanes were isolated, equilibrated in 2x Laemmli buffer (10 min, 37°C), placed on top of SDS-PAGE gels and Western-probed using *anti*-TRPM7 (AB15562, Millipore).

### Complexome profiling

The size distribution of solubilized native TRPM7-associated complexes was investigated using the high-resolution cryo-slicing Blue Native PAGE-mass spectrometry (csBN-MS) technique detailed in [28]. Briefly, membranes isolated from adult mouse brain were solubilized with ComplexioLyte CL-47 (salt replaced by 750 mM aminocaproic acid), concentrated by ultracentrifugation into a 20%/50% sucrose cushion, supplied with 0.125% Coomassie G250 Blue and run overnight on a hyperbolic 1-13% polyacrylamide gel. The region of interest was excised from the lane, proteins fixed in 30% ethanol/15% acetic acid and the gel piece embedded in tissue embedding media (Leica). After careful mounting on a cryo-holder, 0.3 mm slices were harvested, rinsed and subjected to in-gel tryptic digestion as described [28].

### Mass spectrometry

Tryptic digests (dried peptides) were dissolved in 0.5% (v/v) trifluoroacetic acid and loaded onto a C18 PepMap100 precolumn (300 µm i.d. × 5 mm; particle size 5 µm) with 0.05% (v/v) trifluoroacetic acid (5 min 20 µL/min) using split-free UltiMate 3000 RSLCnano HPLCs (Dionex / Thermo Scientific, Germany). Bound peptides were then eluted with an aqueous-organic gradient (eluent A: 0.5% (v/v) acetic acid; eluent B: 0.5% (v/v) acetic acid in 80% (v/v) acetonitrile; times referring to AP-MS/csBN-MS): 5 min 3% B, 60/120 min from 3% B to 30% B, 15 min from 30% B to 99% B or 20 min from 30% B to 50% B and 10 min from 50% B to 99% B, respectively, 5 min 99% B, 5 min from 99% B to 3% B, 15/10 min 3% B (flow rate 300 nL/min). Eluted peptides were separated in a SilicaTip™ emitter (i.d. 75 µm; tip 8 µm; New Objective, USA) manually packed 11 cm (AP-MS) or 23 cm (csBN-MS) with ReproSil-Pur 120 ODS-3 (C18; particle size 3 µm; Dr. Maisch HPLC, Germany) and electrosprayed (2.3 kV; transfer capillary temperature 250/300°C) in positive ion mode into an Orbitrap Elite (AP-MS) or a QExactive HF-X (csBN-MS) mass spectrometer (both Thermo Scientific, Germany). Instrument settings: maximum MS/MS injection time = 400/200 ms; dynamic exclusion duration = 30/60 s; minimum signal/intensity threshold = 2,000/40,000 (counts), top 10/15 precursors fragmented; isolation width = 1.0/1.4 m/z.

Peak lists were extracted from fragment ion spectra using the ’msconvert.exe’ tool (part of ProteoWizard; http://proteowizard.sourceforge.net/; v3.0.6906 for Orbitrap Elite and v3.0.11098 for Q Exactive HF-X; Mascot generic format with filter options ’peakPicking true 1’- and ’threshold count 500 most-intense’). Precursor m/z values were preliminarily searched with 50 ppm peptide mass tolerance, their mass offset corrected by the median m/z offset of all peptides assigned, and afterwards searched with 5 ppm mass tolerance against all mouse, rat, and human (mouse/rat brain samples) or only human (HEK293 cell samples) entries of the UniProtKB/Swiss-Prot database (release 20190731 / 20200226, respectively). Acetyl (Protein N-term), Carbamidomethyl (C), Gln->pyro-Glu (N-term Q), Glu->pyro-Glu (N-term E), Oxidation (M), Phospho (S, T, Y), and Propionamide (C) were chosen as variable modifications, and fragment mass tolerance was set to ±0.8 Da (Orbitrap Elite data) or ± 20 mmu (Q Exactive HF-X data). One missed tryptic cleavage was allowed. The expect value cut- off for peptide assignment was set to 0.5. Related identified proteins (subset or species homologs) were grouped using the name of the predominant member. Proteins either representing exogenous contaminations (e.g., keratins, trypsin, IgG chains) or identified by only one specific peptide were not considered.

Label-free quantification of proteins was carried out as described in (36, 60). Peptide signal intensities (peak volumes, PVs) from FT full scans were determined, and offline mass calibrated using MaxQuant v1.6.3 (http://www.maxquant.org). Then, peptide PV elution times were pairwise aligned using LOESS regression (reference times dynamically calculated from the median peptide elution times overall aligned datasets). Finally, PVs were assigned to peptides based on their m/z and elution time (±1 min / 2-3 ppm, as obtained directly or indirectly from MS/MS-based identification). PV tables were then used to calculate protein abundance ratios in AP versus control (Figure 1C, see below), the abundance norm value (Figure 1B, lower right) as an estimate for molecular abundance (both described in (33)), and csBN-MS abundance profiles (Figure 1B, lower left) as detailed in (60). The latter were smoothed using a sliding average calculated over a window of 5 slices. Slice numbers were converted to apparent complex molecular weights by the sigmoidal fitting of log(MW) versus slice number of the observed profile peak maximum of mitochondrial marker protein complexes (61). For identification of proteins specifically and consistently co-purified with TRPM7 (Figure 1C, table 1), data obtained from twenty-one TRPM7 affinity-purifications (mouse brain: four antibodies, two solubilisation conditions; rat brain: three antibodies, two solubilisation conditions; HEK293 cells: four antibodies, one or two solubilisation conditions) with their corresponding controls (APs with four or five different target-unrelated control (TUC) antibodies as negative controls for *anti*-TRPM7 APs from rodent brain; *anti*-TRPM7 APs from TRPM7^-/-^ HEK293 cells (12) as negative controls for *anti*-TRPM7 APs from WT HEK293 cells; anti-HA AP from HEK293 cells heterologously expressing TRPM7-myc as negative control for *anti*-HA AP from HEK293 cells overexpressing TRPM7-HA) were evaluated using the Belki software suite (https://github.com/phys2/belki). Thus, protein abundance ratios from anti- TRPM7 APs versus controls were normalized to the background and the abundance ratio of TRPM7, resulting in so-called protein ratio distance values (RDs, see supplementary file 1 to figure 1). The latter were then visualized and the proteins with the highest (effective cutoff 0.25) and most consistent (at least two of the three or four APs) RDs were then manually inspected for the quality of the underlying LC-MS data (i.e. the assignment, internal consistency and signal-to-noise of the respective PV values).

### Heterologous expression of TRPM7, CNNM3 and ARL15 in *Xenopus laevis* oocytes

#### Two-electrode voltage clamp (TEVC) measurements

TEVC measurements were performed as described previously (6) with a few modifications. Linearized cDNAs of TRPM7 (in pOGI), TRPV1 (in pNKS2), CNNM3 and ARL15 (both in pCMV6-Entry) were used for *in vitro* synthesis of cRNA (T7 or SP6 mMESSAGE mMACHINE™ transcription kits (Thermo Fisher Scientific). In Figure 4A, oocytes were injected with 5 ng of TRPM7 cRNA or co-injected with 2.5 ng of CNNM3 (2:1 ratio), 2.5 ng Arl15 (2:1 ratio), and 2.5 ng of CNNM3 with 2.5 ng of Arl15 cRNAs (2:1:1 ratio). In Figure 4B, oocytes were co-injected with 5 ng of TRPM7 and 0.025-0.5 ng of Arl15 cRNAs (200:1- 10:1 ratio).

The injected oocytes were kept in ND96 solution (96 mM NaCl, 2 mM KCl, 1mM CaCl_2_, 1 mM MgCl_2_, 5 mM HEPES, pH 7.4), supplemented with 20 μg/ml gentamicin at 16°C. TEVC measurements were performed three days after injection at room temperature in Ca^2+^/Mg^2+^-free ND96 containing 3.0 mM BaCl_2_ instead of CaCl_2_ and MgCl_2_ using a TURBO TEC-05X amplifier (npi electronic) and CellWorks software (npi electronic). Oocytes were clamped at a holding potential of −60 mV, and 0.5 ms ramps from −80 mV to +80 mV were applied at 6 s intervals. For statistical analysis, outward current amplitudes were extracted at +80 mV for individual oocytes. Statistical significance (ANOVA) was calculated using GraphPad Prism 7.03.

#### Western blot

Oocytes (n=6 per group) were treated with a lysis buffer (Pierce IP Lysis Buffer, Pierce) containing protease inhibitor and phosphatase inhibitor cocktails (Biotool), mixed (1:1) with 2x *Laemmli* buffer, heated at 70°C for 10 min, and cooled on ice. Samples were separated by SDS-PAGE (4-15% gradient Mini-PROTEAN, Bio-Rad) and electroblotted on nitrocellulose membranes (GE Healthcare Life Science). After blocking with 5% (w/v) non-fat dry milk in Tris-buffered saline with 0.1% Tween 20 (TBST), the upper part of the membrane was incubated with *anti*-M7d antibody (0.2 µg/ml) diluted in TBST with 5% (w/v) BSA, followed by washing in TBST, incubation with a horseradish peroxidase-coupled anti-mouse lgG (Cell Signaling Technology; 1:1000 in TBST with 5% (w/v) non-fat dry milk), and washing again in TBST. Blots were visualised using a luminescence imager (ChemiDoc imaging System, Bio-Rad). The lower part of the membrane was developed using an *anti*-Na^+^/K^+^ ATPase antibody (EP18459, Abcam; 1:1000).

#### Immunofluorescent staining

Oocytes were fixed in 4% (w/v) PFA (Electron Microscopy Sciences) in ND96 solution for 15 min at room temperature (RT), followed by incubation in ice-cold methanol for 60 min at -18°C. After washing in ND96 (3x, RT), oocytes were incubated in ND96 containing 5% (w/v) BSA for 30 min at RT. *Anti*-M7d antibody (1.6 µg/ml in ND96 with 5% BSA) was applied overnight at 4°C. Afterwards, oocytes were washed in ND96 (3x, RT), and a goat *anti*-mouse antibody conjugated with Alexa Fluor 488 (Life Technologies; 2 μg/ml in ND96 with 5% BSA) was applied for 1h at RT. After washing in ND96 (3x, RT), differential interference contrast (DIC) and confocal images were obtained with a confocal laser scanning microscope LSM 880 AxioObserver (Carl Zeiss). We used a Plan-Apochromat 10x/0.45 objective, 488 nm excitation wavelengths and 493–630 nm filters. Acquired DIC and confocal images were analysed using the ZEN2.3 software (Carl Zeiss).

## Acknowledgements

V.C., T.G., U.S. and B.F. were supported by the Deutsche Forschungsgemeinschaft (German Research Foundation, DFG), TRR 152 (P02 and P15). B.F. and U.S. were supported by the DFG under Germany’s Excellence Strategy (CIBSS-EXC2189 project ID: 390939984) and Project-ID 403222702 – SFB 1381. A.N. was supported by the DFG Project-ID 335447717 – SFB 1328 (P15). We thank Veit Flockerzi for anti- TRPC1/3 antibodies and David Clapham for TRPM7-/- HEK293 cells. We thank Joanna Zaisserer and Anna Erbacher for their technical assistance.

## Author Contributions

Funding Acquisition, B.F., U.S., V.C. and T.G.; Investigation, A.K., F.T.T., C.M., W.B., and V.C.; Formal Analysis, A.H., A.N., U.S.; Writing – Original Draft, V.C. and T.G.; Writing – Review & Editing, A.K., A.N., U.S. and B.F.

## Supplementary figure legends

**Figure 1-figure supplement 1.**
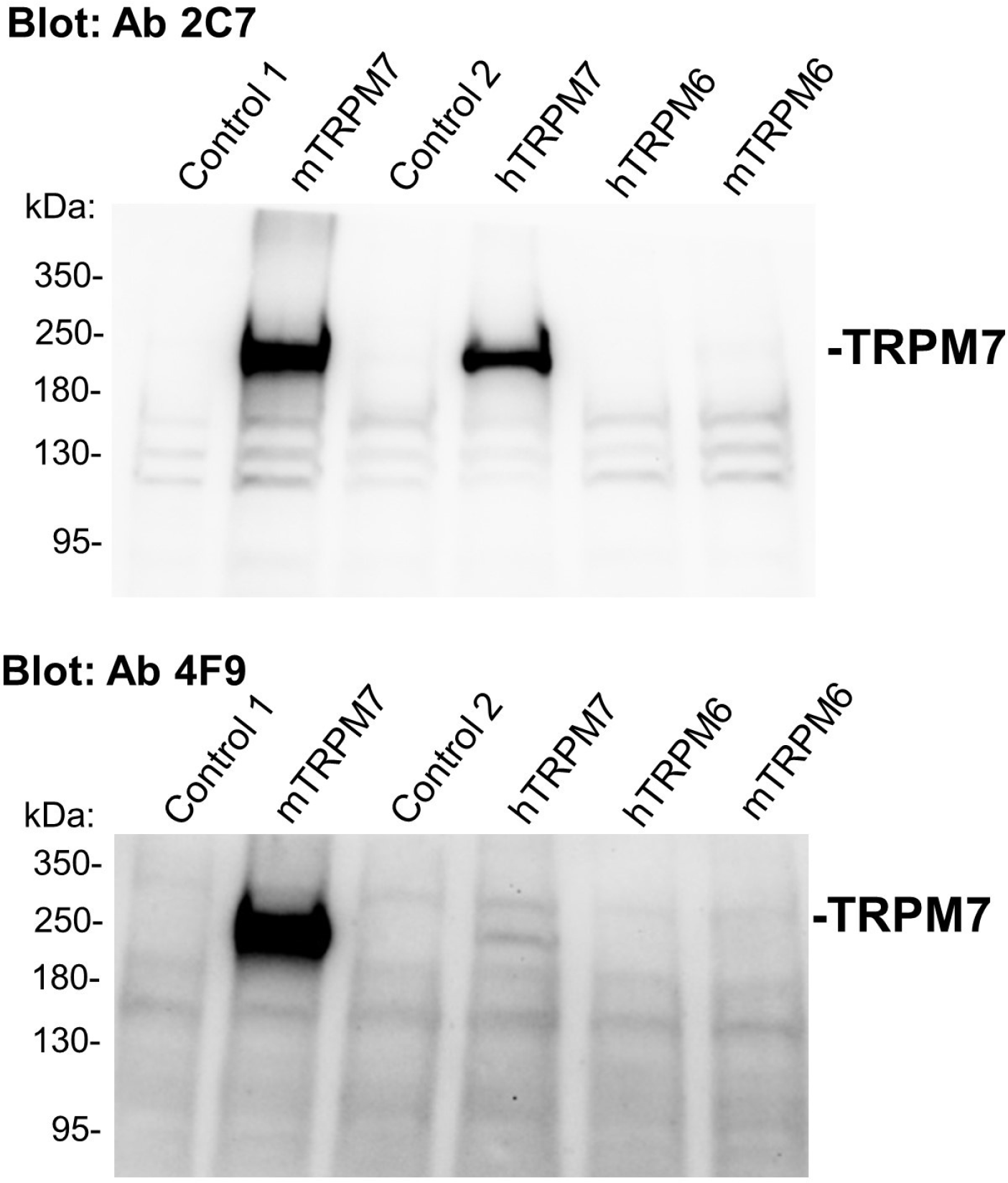
The specificity of an *anti-*TRPM7 mouse monoclonal antibody in Western blot assessment of the recombinant TRPM6 and TRPM7 proteins. 0.8 µg/ml of 2C7 IgG (*Upper panel*) or 1.3 µg/ml of 4F9 IgG (*Lower panel*) were used for Western-blot analysis of untransfected HEK293 cells (*Control 1*) or cells transfected with mouse TRPM7 cDNA *(mTRPM7)*, uninducted *(Control 2)* or induced HEK293 T-Rex cells expressing the human TRPM7 protein (*hTRPM7)*, HEK293 cells transfected with human *(hTRPM6*) or mouse TRPM6 cDNA *(mTRPM6)*. Red arrows indicate the expected location of the TRPM7 band. Representative results of two independent experiments are shown.

**Figure 3-figure supplement 1.**
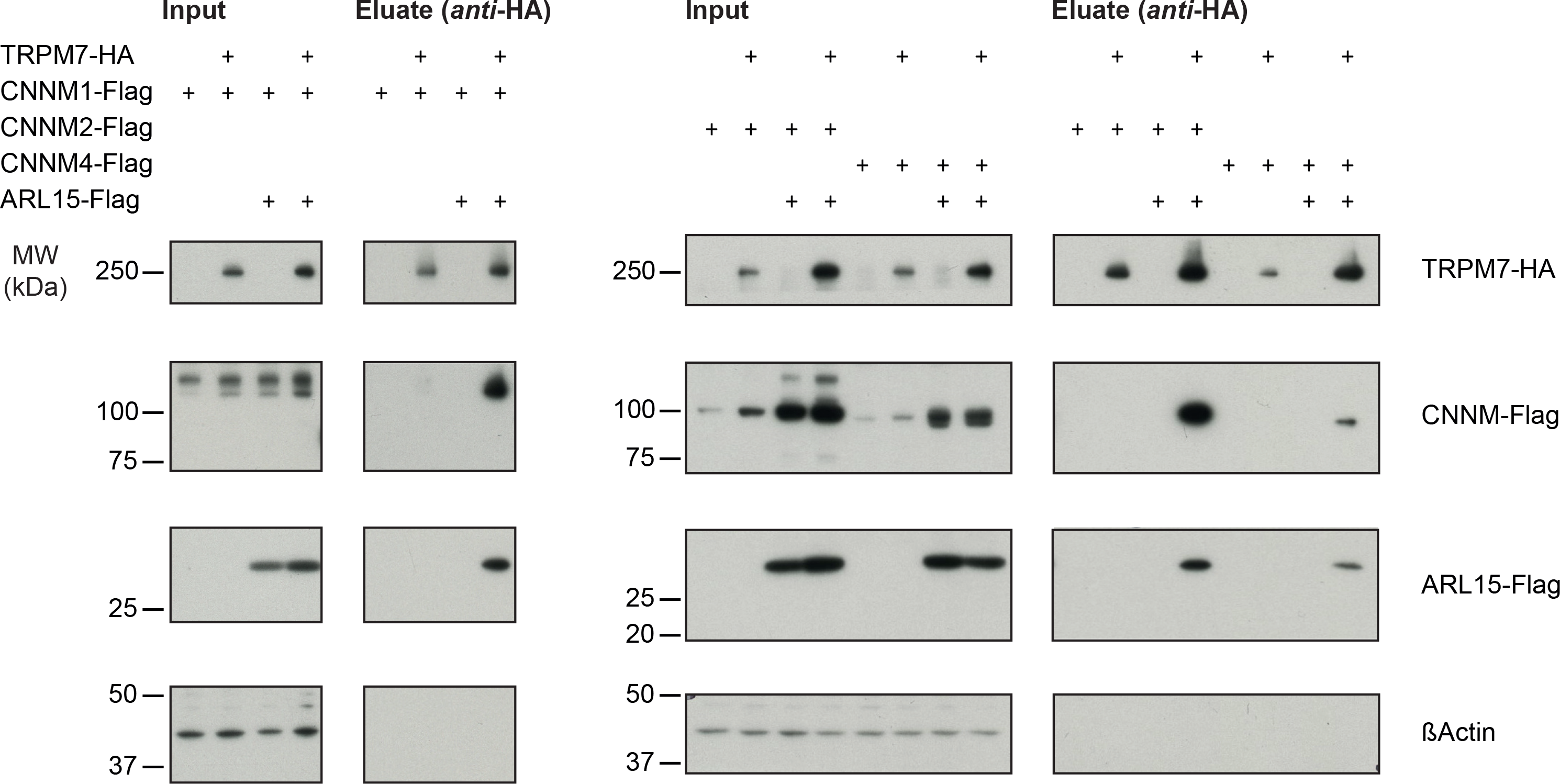
Heterologous reconstitution of TRPM7 complexes in HEK293 cells. APs with *anti*-HA antibody from CL-47 solubilised membrane fractions of TRPM7^-/-^ HEK293 cells transiently expressing the indicated combinations of proteins. Input and eluates of the distinct APs were separated by SDS-PAGE and Western-probed with anti-Flag and *anti*-HA antibodies. MW is marked on the left. Robustness of this pulldown assay was verified with reconstituted TRPM7/CNNM3**/**ARL15 complex (figure 3, three replicates) before CNNMs1, 2 and 4 were confirmed each in one separate reconstitution experiment.

**Figure 4-figure supplement 1.**
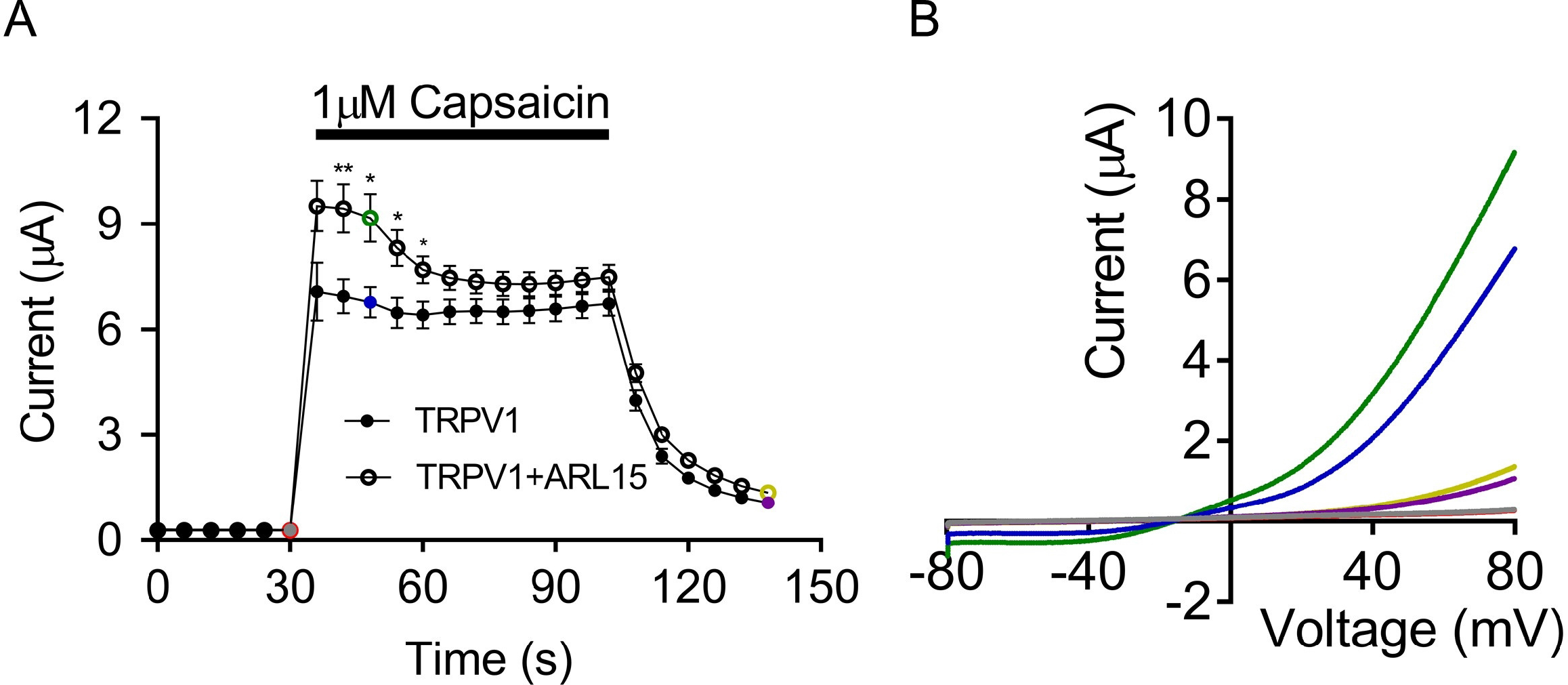
Two-electrode voltage clamp (TEVC) measurements of capsaicin-induced TRPV1 currents. Voltage ramps from -80 to +80 mV were applied every 5 s, and current amplitudes (mean ± SEM, n=7) were acquired at +80 mV in *Xenopus* oocytes expressing TRPV1 alone or TRPV1 with ARL15 (cRNA ratio 2:1) and plotted over time. Oocytes were perfused with 1 µM capsaicin as indicated by the black bar. **(A)** Representative current-voltage (I-V) relationships of TRPV1 currents shown in (A) prior, during and after exposure of oocytes to capsaicin as indicated by the corresponding coloured data points.

## Supplementary files

**Supplementary file 1 to** **Figure 1** **and** Table 1. Numerical data for peak volumes, abundance norm values, relative abundance and ratio distance values obtained through analysis of the MS data.

<u>Excel file</u> contains twenty-two worksheets.

**Supplementary file 2 to** **Figure 2**. Phosphorylation sites in TRPM7, CNNM3 and CNNM4 identified in APs from HEK293 cells and rodent brain.

<u>Excel file</u> contains one worksheet:

The phosphorylated residues of TRPM7, CNNM3 and CNNM4 identified by MS in the present study are outlined in conjunction with previously published data (62-65).

**Supplementary file 3 to** **Figure 2**. MS-spectra of phosphorylated TRPM7, CNNM3, and CNNM4 peptides identified in APs from HEK293 and rodent brain.

<u>Word file</u>

## Data availability

The mass spectrometry proteomics data have been deposited to the ProteomeXchange Consortium via the PRIDE (66) partner repository with the dataset identifier PXD025279 and 10.6019/PXD025279.

## Reviewer account details

Username: Password: yi6BamNW

